# Prophage ΦESI promotes competitive dominance in the emergent *Salmonella* serovar Infantis lineage

**DOI:** 10.64898/2026.03.17.712176

**Authors:** Cristobal Martinez-Padilla, Daniel Tichy-Navarro, Rocio Barron-Montenegro, Ignacia Herrera-Vergini, Juan A. Ugalde, Diana Álvarez-Espejo, Andrea I. Moreno-Switt, Alejandro Piña-Iturbe

**Author notes:** Correspondence: Alejandro Piña-Iturbe; Andrea I. Moreno-Switt.

## Abstract

The global spread of multidrug-resistant *Salmonella enterica* serovar Infantis has been largely attributed to the pESI megaplasmid, yet additional factors underlying the ecological success of this lineage remain unclear. Here, we characterize ΦESI (<u>ph</u>age of <u>e</u>mergent *<u>S</u>almonella* Infantis), a temperate bacteriophage discovered during coculture of an emergent pESI-positive strain (PM57) and a non-emergent strain (DR006). Phage ΦESI shows a siphovirus morphotype with a long tail and an elongated head, with a genome of 46,490 bp, which is integrated as a prophage in the 3’-end of an Arg-tRNA gene in PM57, but is absent in DR006, which is susceptible to the ΦESI-mediated lysis. The screening of ΦESI genes across 20,429 global *Salmonella* Infantis genomes revealed an almost exclusive association of ΦESI and ΦESI-like phages with the emergent pESI-positive *Salmonella* Infantis lineage. Further bioinformatic analyses of complete chromosomes revealed diverse ΦESI insertion profiles showing geographic clustering, and the presence of large-scale chromosomal inversions flanked by ΦESI genes. Competition assays showed that PM57 outcompeted DR006 in coculture, coinciding with high viral loads that selectively targeted DR006, and susceptibility assays showed that strains lacking ΦESI/ΦESI-like prophages were susceptible to ΦESI-mediated lysis. Together, our findings identify ΦESI as a competitive factor associated with emergent *Salmonella* Infantis, able to selectively eliminate susceptible competitors. Our findings suggest that ΦESI/ΦESI-like prophages contributed to the persistence and global dissemination of the emergent *Salmonella* Infantis lineage.

**Importance:** Emergent multidrug-resistant *Salmonella enterica* serovar Infantis strains carrying pESI megaplasmids have spread worldwide, posing a global public health threat and a significant economic burden. Nevertheless, the factors contributing to the success of this foodborne pathogen, beyond pESI, remain poorly understood. Here, we describe ΦESI, a novel temperate bacteriophage carried by the emergent lineage as a chromosomally integrated prophage. We show that ΦESI provides a competitive advantage to emergent strains by selectively killing non-emergent competitors while protecting lysogens from reinfection. Our findings uncover the contribution of ΦESI/ΦESI-like bacteriophages to the success of emergent *Salmonella* Infantis, highlighting how lineage-associated prophages can shape the ecological success of pathogenic bacteria.

## INTRODUCTION

Non-typhoidal *Salmonella* (NTS) is a major cause of foodborne disease worldwide, affecting millions of people and causing approximately 215,000 deaths annually (1). Usually producing self-limiting gastroenteritis, NTS can also cause a severe invasive infection requiring antibiotic treatment, including the use of third-generation cephalosporins, fluoroquinolones, and azithromycin (2). While more than 2,600 *Salmonella* serovars have been identified, most human cases are caused by a limited number of highly prevalent serovars (3–5).

Over the past two decades, multidrug-resistant (MDR) and extended-spectrum β-lactamase (ESBL)-producing *Salmonella* enterica serovar Infantis have emerged as a global zoonotic pathogen primarily associated with consumption of chicken meat, likely shaped by international poultry trading (6, 7). The success of emergent *Salmonella* Infantis is mainly linked to the acquisition of a ≈300 Kbp pESI/pESI-like megaplasmid carrying multiple antimicrobial resistance genes (e.g. *aph(3’)-Ia*, *aph*(*4*)*-Ia*, *aac*(*3*)*-Iva*, *floR*, *fosA3*, *sul1*, *tet(*A), *bla*_CTX-M-1_, *bla*_CTX-M-65_) along with virulence (e.g. yersiniabactin synthesis locus, K88-like fimbria, Infantis plasmid-encoded fimbria), disinfectant-resistance (*qacE*Δ*1*), and heavy-metal tolerance (e.g. *mer* operon) determinants (8–13). Moreover, chromosomal mutations in the quinolone resistance-determining region (QRDR) of the *gyrA* gene are common in the emergent *Salmonella* Infantis lineage (7). Together, these acquired features contribute to enhanced chicken gut colonization, increased shedding, persistence in chicken meat processing facilities, and resistance to multiple antibiotics including those used in chicken farms, and as first-line treatment for severe infections in humans (8, 9, 14–17). In addition, the widespread dissemination of emergent *Salmonella* Infantis across all continents (except Antarctica) and among multiple environments and niches, including surface/irrigation waters, wild animals, livestock, companion animals, poultry, and food, makes this MDR pathogen a major threat to global public health (7, 18–20). While strong evidence supports pESI-like megaplasmids as major contributors to the successful dissemination of emergent *Salmonella* Infantis, the impact or potential role of other genetic factors remains unexplored.

Bacteriophages (phages), highly diverse viruses that infect bacteria, are major contributors to bacterial diversification, shaping their genetic content and ecological interaction (21). They replicate primarily through two cycles: the lytic cycle, in which active phage replication leads to the production, assembly, and release of new viral particles, lysing the host cell (22), and the lysogenic cycle, in which the phage genome integrates into the bacterial chromosome as a prophage and is passively inherited during cell division (23). Phages that undergo only the lytic cycle are termed virulent or strictly lytic phages, while temperate phages can follow either the lytic or lysogenic pathway (23). During lysogenic infection, the temperate phage genome integrates into the bacterial chromosome either randomly or at specific “attachment sites” (*att* sites), through a reaction catalyzed by integrases encoded by the phage genome (24, 25). The lysogenic state is maintained by transcriptional repressors encoded by lysogenic master regulator genes within the prophage genome; these repressors inhibit the expression of lytic genes such as holins and lysins (24). Prophages can be reactivated through induction by SOS-dependent or SOS-independent mechanisms, including exposure to DNA-damaging agents (e.g. UV radiation), reactive oxygen species, quorum-sensing signals, phage-encoded regulatory systems, phage–phage interactions, or spontaneous induction (24, 26–30).

Prophages are common in clinically and environmentally important bacteria, and they dominate the virome in the gut-associated microbiomes (31–33). They influence community structure through lysogeny, lysis, and horizontal gene transfer, enhancing bacterial genetic flexibility and adaptation (33, 34). Prophages can also provide competitive advantages to their bacterial hosts against susceptible subpopulations. For example, the Lambda prophage can benefit its *Escherichia coli* hosts, acting as a “replicative toxin” against susceptible bacteria, improving the survival of its carriers in both solid and liquid environments (35). Similarly, *Enterococcus faecalis* V583 uses phage particles to maintain dominance in its intestinal niche (36). Moreover, in *Salmonella* Typhimurium, spontaneous induction of the Gifsy prophage provides a strong growth advantage in coculture, underscoring how prophage activation can decisively influence bacterial competition (37).

Only a few comparative studies have assessed the prophage content of *Salmonella* Infantis among emergent and non-emergent isolates. Although these works were limited by relatively small genome datasets, they showed evidence of specific prophages that correlate with phylogenetic clades of *Salmonella* Infantis and were unique to emergent genomes (38, 39), suggesting a potential role of these prophages in the diversification of this foodborne pathogen. In this study, we describe the isolation and characterization of ΦESI (phage of <u>e</u>mergent *<u>S</u>almonella* Infantis), a temperate bacteriophage associated with the emergent *Salmonella* Infantis lineage, providing compelling evidence that it confers a competitive advantage over non-emergent strains lacking this prophage.

## RESULTS

### **Φ**ESI is a lysogenic bacteriophage integrated into the chromosome of pESI-positive *Salmonella* Infantis PM57

The bacteriophage ΦESI was isolated from a coculture of *Salmonella* Infantis strain PM57 (pESI-positive; emergent lineage) and strain DR006 (pESI-negative; non-emergent lineage) **(Fig. 1A)**. ΦESI produced clear lytic plaques approximately 1 mm in diameter on double-layer agar plates using DR006 as the host strain **(Fig. 1B, Fig. S1)**. Transmission electron microscopy revealed that ΦESI exhibits a siphovirus morphotype, characterized by a long, flexible tail (165.6 ± 5.7 nm) and an elongated head structure (length: 89.2 ± 4.9 nm; width: 39.6 ± 4.9 nm) **(Fig. 1C, Fig. S2, Table S1)**. The genomic analysis showed that ΦESI has a 46,490 bp genome with a 49.3% G+C content and 88 CDSs according to the Pharokka annotation. The genome was flanked by 127 bp-long direct terminal repeats and was predicted to encode several lysogeny-associated proteins, including an integrase, suggesting a lysogenic lifestyle. **(Fig. 1D; Table S2)**. A BLASTn search of the NCBI Virus nucleotide database using the genome of ΦESI yielded hits with ≤51% coverage. The two phage genomes showing the highest nucleotide identity had coverage values of 24% (phage Inf 5-2, 99% identity; GenBank accession OR850760.1) and 51% (phage FSL SP-016, 97% identity; GenBank accession KC139516.1). These results together with the taxonomic analyses performed using VICTOR, VIRIDIC and the top eight phage genomes from the NCBI Virus search, indicated that ΦESI may represent a novel bacteriophage genus within the class *Caudoviricetes* (**Fig. S3**).

**Figure 1.**
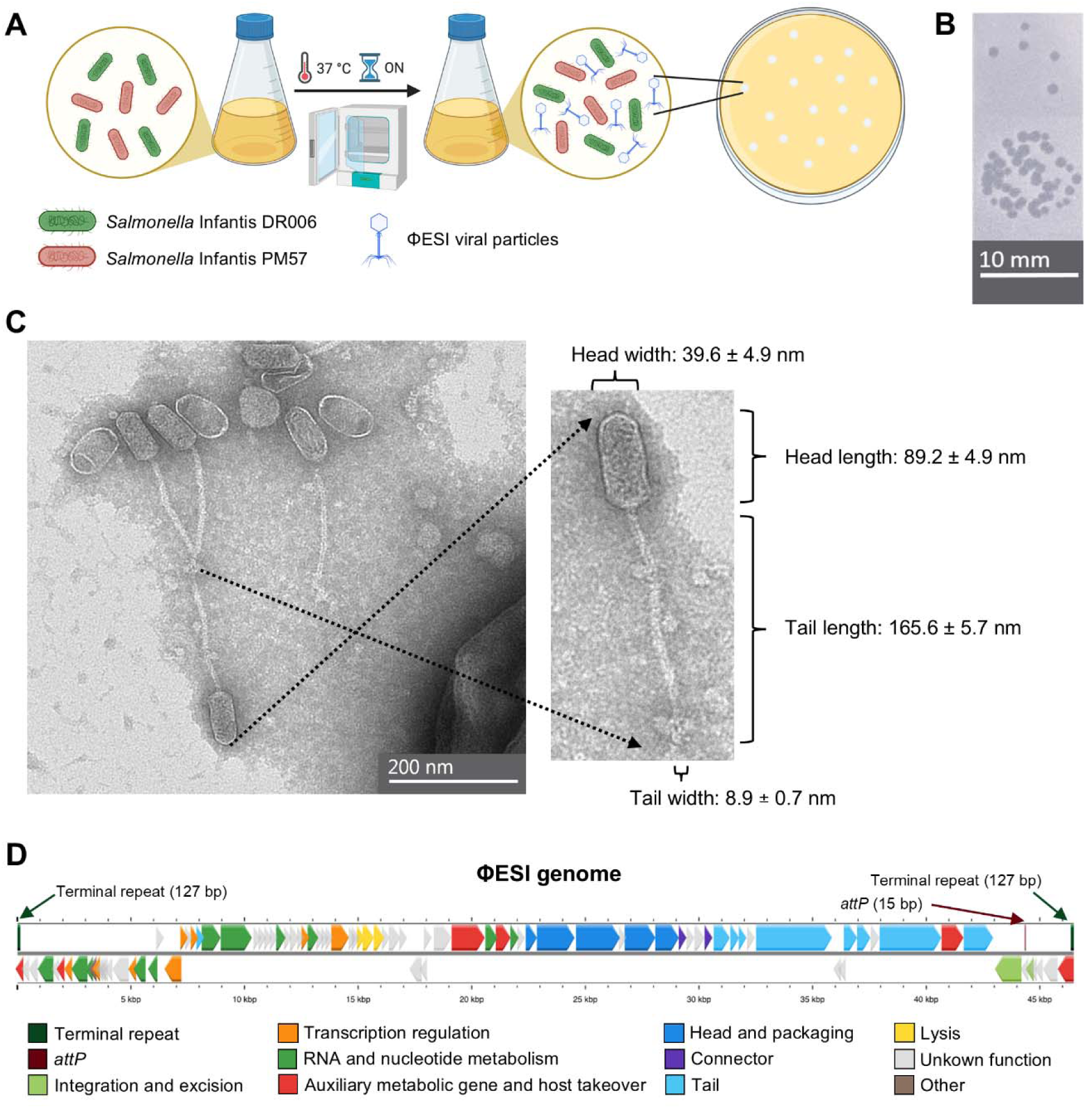
Isolation, Phenotypic and Genotypic Characterization of ΦESI. **A)** Schematic representation of the experimental procedure used to isolate phage ΦESI where *Salmonella* Infantis PM57 (red), from the emergent lineage, and the non-emergent isolate DR006 (green) were cocultured (ON: overnight) on trypticase soy broth media, and the filtered supernatant used for phage isolation. **B)** Lysis plaque morphology of ΦESI on a double-layer agar of *Salmonella* Infantis DR006. A bar scale of 10 mm is shown. **(C)** Transmission electron micrograph of the ΦESI bacteriophage. The tail and head length and width (mean ± standard deviation) were estimated from measurements of seven viral particles. **D)** Lineal representation of the ΦESI phage genome. Genes were colored according to their predicted function as determined by the Pharokka annotation. The 127-bp terminal repeats and the 15-bp *attP* site are indicated with arrows.

To assess the source of ΦESI, BLASTn alignments were carried out against the genomes of the two cocultured strains, PM57 and DR006. The analysis revealed the presence of ΦESI as a prophage integrated within the chromosome of strain PM57 at the 3’-end of an Arg-tRNA-encoding gene, spanning positions 1,451,172 to 1,497,549 **(Fig. 2)**, while it was absent from the genome of strain DR006 **(Fig. 2A)**. The BLAST results also revealed that the integrated prophage, compared with the virion genome, presented a translocation of 2,138 bp that placed the region located after the integrase gene at the beginning of the prophage genome (**Fig. 2B**). In addition, the 127bp-long terminal repeats were combined in a single, internal, copy in the prophage, which was instead flanked by 15bp-long attachment sites (*attL* and *attR*) that shared full nucleotide identity with the last nucleotides of the Arg-tRNA gene, which is not occupied in DR006. Consistent with these findings, ΦESI formed lytic plaques on DR006, whereas the strain PM57 appeared resistant to plaque formation (**Fig. S4**). Our analyses confirmed that the isolated bacteriophage came from the genome of strain PM57.

**Figure 2.**
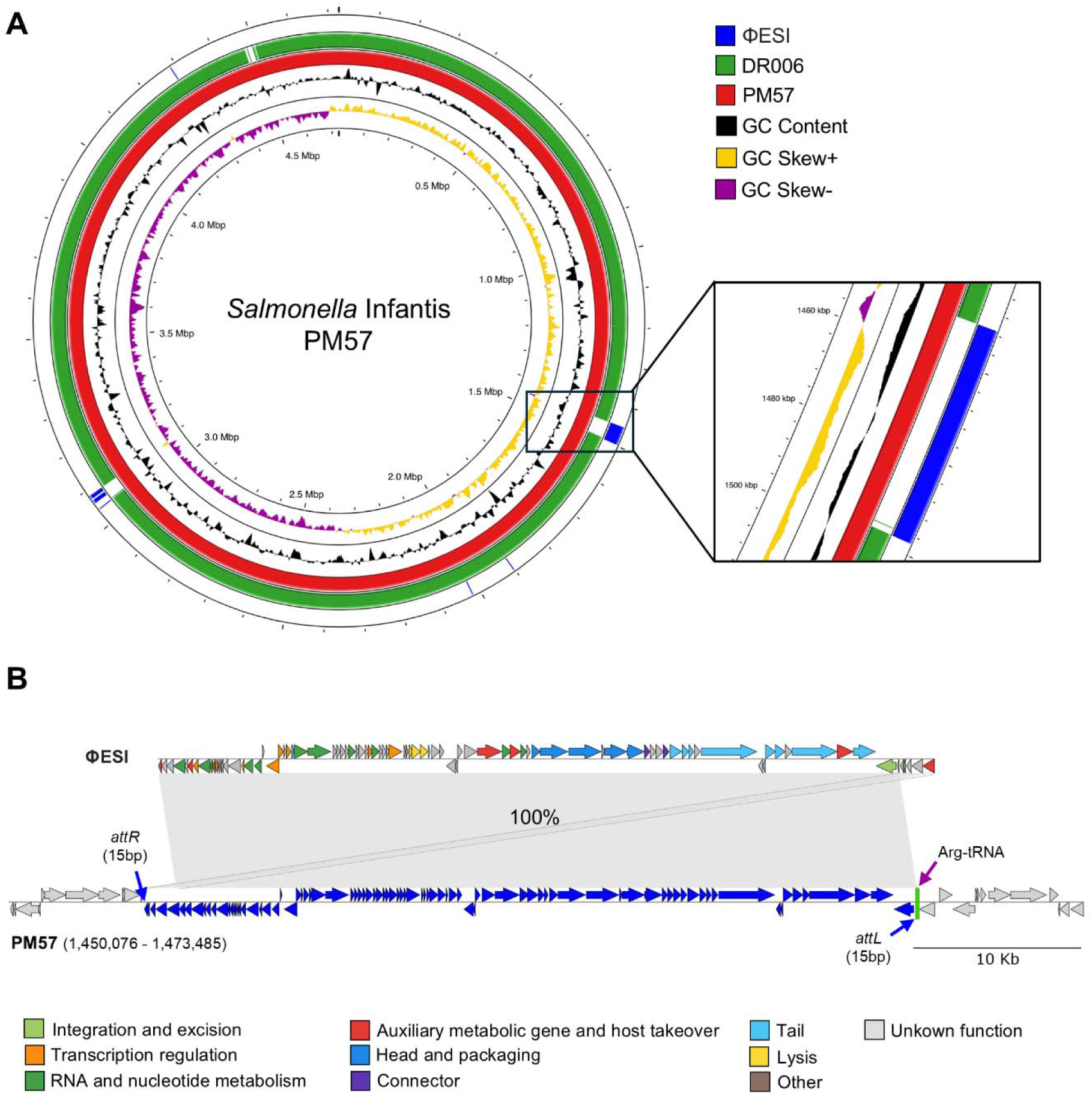
ΦESI is integrated as a prophage in the chromosome of *Salmonella* Infantis PM57. **A)** Circular representation of the complete *Salmonella* Infantis PM57 chromosome (red) and the BLAST alignments of DR006 (green), and the genome of bacteriophage ΦESI (blue). A zoom-in view of the region where ΦESI is integrated in the PM57 chromosome is also presented. **B)** BLAST alignment between the bacteriophage ΦESI genome and the prophage ΦESI region integrated into chromosome of PM57 strain. Genes were colored according to their predicted function as determined by the Pharokka annotation using the same color palette as in Fig. 1D. The insertion site at the Arg-tRNA and the flanking 15-bp attachment sites are indicated with purple and blue arrows.

### **Φ**ESI and **Φ**ESI-like prophages are associated with emergent *Salmonella* Infantis and major chromosome rearrangements

To assess the global distribution of the ΦESI prophage across *Salmonella* Infantis, we conducted bioinformatic and phylogenetic analyses based on 36,499 genomes publicly available in Enterobase (64) as of September 9^th^, 2025. A search of the 88 ΦESI CDSs among 15,707 non-clonal genomes revealed a variable conservation of these CDSs, revealing four major groups based on the frequency of CDSs present (**Fig. 3A, B; Tables S3, S4**). The first group represented a 39.9% (6,272/15,707) and carried from 86 to 88 ΦESI CDSs; the second group represented the 15.3% (2,410/15,707) and carried 73 CDSs; the third group represented the 17.4% (2,726/15,707) and carried 16-40 CDSs; and the fourth group represented the 16.4% (2,580/15,707), carrying zero ΦESI CDSs. The remaining 1,719 genomes (10.9%) represented low frequency cases carrying an intermediate number of CDSs between the four major groups. Based on this distribution, we classified the genomes in three categories: ΦESI-positive, defined as genomes containing more than 86 phage CDSs; ΦESI-like-positive, defined as genomes containing 73 to 85 phage CDSs; and ΦESI-negative genomes, harboring fewer than 73 ΦESI CDSs. According to this grouping, a statistically significant association (χ², p < 0.0001) was found between the presence of ΦESI and ΦESI-like prophages and the emergent lineage of *Salmonella Infantis*, represented by the carriage of pESI-like megaplasmids. Specifically, 92.5% (n = 8,978/9,706) of emergent genomes harbored the prophages compared with only 7.7% (n = 464/6,001) of non-emergent genomes (**Fig. 3C, D**). The odds ratio computed from the contingency table was 147.2 (95% CI 130.4-166.1), indicating that plasmid-carrying genomes had ∼147-fold higher odds of prophage carriage than genomes lacking the plasmid. Together, these findings suggest that ΦESI-like prophages are tightly linked to the globally emergent pESI-positive *Salmonella Infantis* lineage.

**Figure 3.**
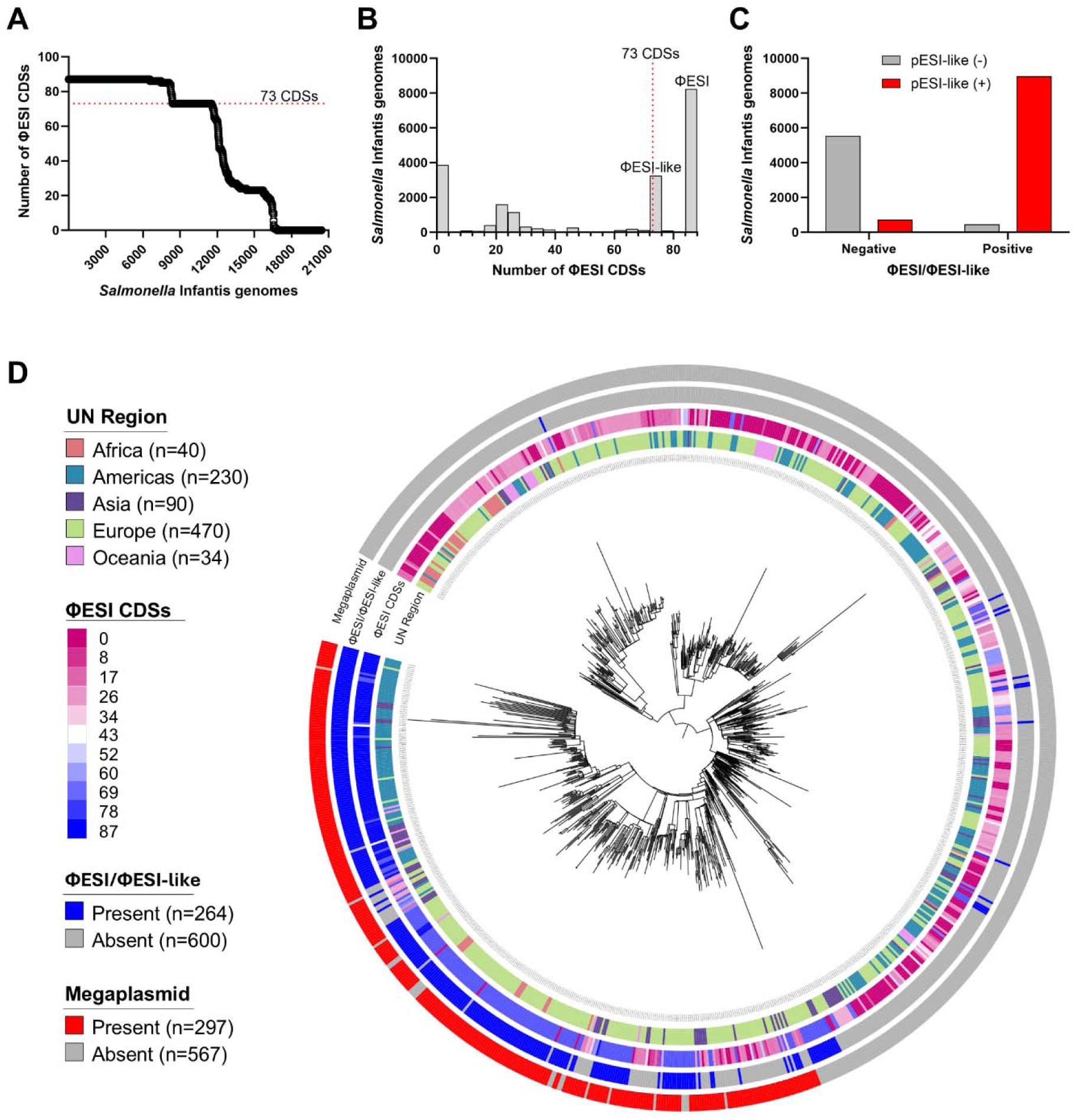
ΦESI and ΦESI-like prophages are associated with pESI-positive *Salmonella* Infantis. **A)** Number of the ΦESI CDSs found across the global dataset of 20,429 *Salmonella* Infantis genomes. **B)** Histogram showing the distribution of the global genome dataset based on the number of ΦESI CDSs. The red broken line in A and B indicate the cut-off value used to consider a genome positive for ΦESI or ΦESI-like prophages. **C)** Frequency of pESI-positive or - negative *Salmonella* Infantis genomes in the global dataset categorized by the presence of ΦESI or ΦESI-like prophages. **D)** Recombination-free core-SNP maximum likelihood phylogeny of 864 *Salmonella* Infantis genomes representing the global geographic, temporal and genomic diversity of *Salmonella* Infantis from the HC200_36 cluster. The tree is rooted at the best-fitting temporal root. The rings show, from the inside to the outside, the geographic origin of the genomes at the continental (UN region) level, the number of ΦESI CDSs found in those genomes, the presence of pESI-like megaplasmids, and the presence of ΦESI and ΦESI-like prophages based on the ≥73-CDS cut-off.

To assess whether this strong association is accompanied by conserved chromosomal integration patterns, we screened 146 closed *Salmonella* Infantis chromosomes available in the NCBI Pathogen Detection database (**Table S5**) for the presence of the complete ΦESI genome sequence. To enable position-based comparisons, chromosome sequences were rearranged, if needed, to start at the *dnaN* gene. Our screening detected ΦESI-related sequences (≥5% coverage and ≥80% identity) in 105/146 (71.9%) genomes, all of which were pESI-positive, exhibiting different completeness patterns (**Table S6**). Notably, the sequences were found in two conserved insertion loci located at positions ≈1.45 Mbp and ≈3.05 Mbp, delimiting a ≈1.5 Mbp intermediate region (**Fig. 4A**). Overall, 25/105 ΦESI-positive genomes (23.8%) contained ΦESI-related sequence at or near the 1.45 Mbp locus, and 80/105 (76.2%) displayed hits at both loci. No genome was found with ΦESI hits only at the 3.05 Mbp. We also detected that, for 39.0% (41/105) genomes, the ΦESI insertion loci co-localized with the boundaries of a large-scale chromosomal inversion of the 1.5 Mbp intermediate region (**Fig. 4B**) in comparison to the PM57 strain.

**Figure 4.**
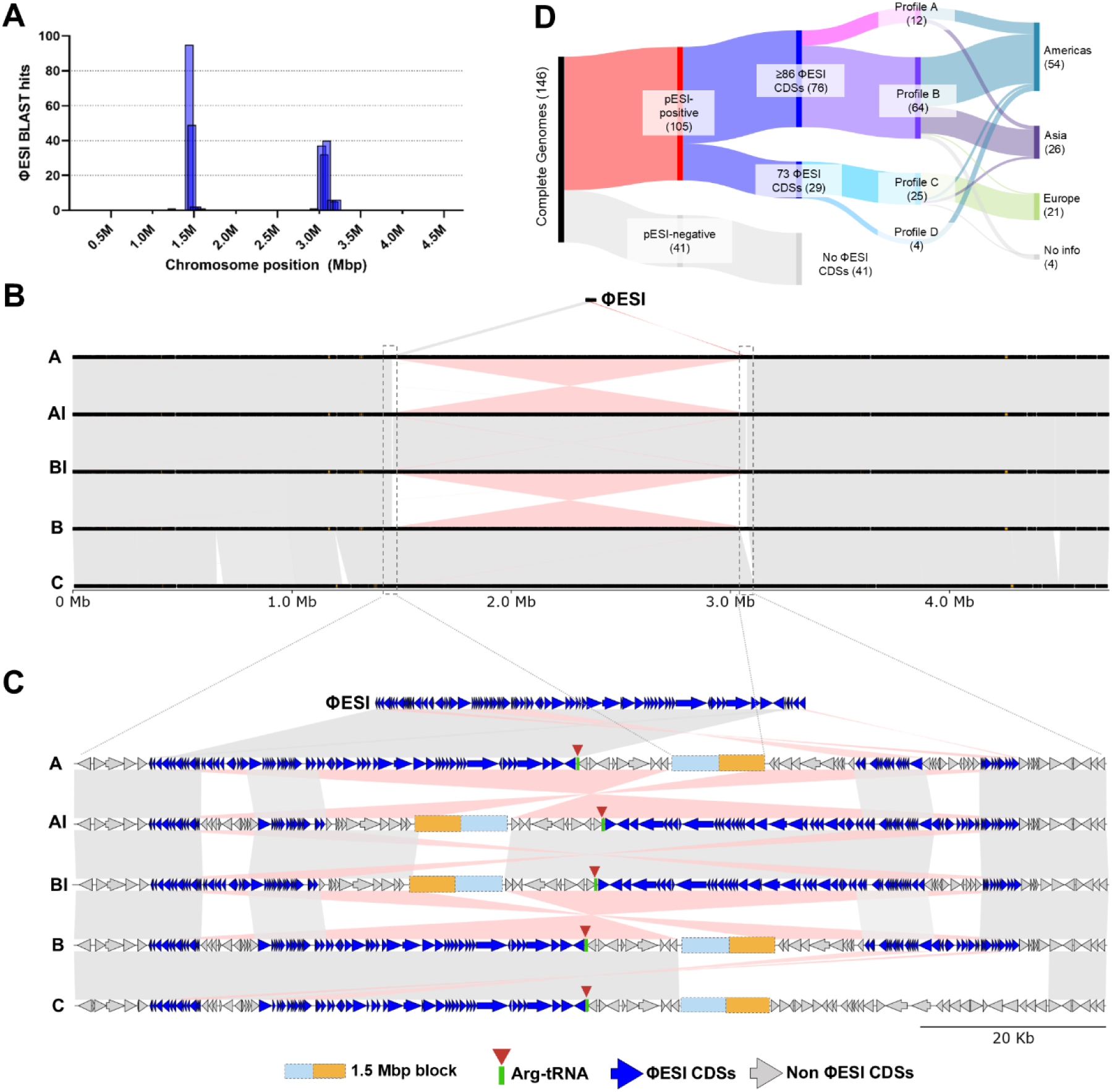
ΦESI and ΦESI-like prophages are found at conserved chromosome locations and are related to large chromosomal inversions. **A)** Histogram showing the distribution of ΦESI-derived ABRicate hits across chromosomal coordinates among 105 *Salmonella* Infantis closed genomes. **B)** BLAST alignment of the chromosome of representative isolates with different profiles of ΦESI insertion showing a large-scale chromosomal inversion flanked by the ΦESI/-like insertion sites. BLAST hits (>99% identity) are represented as grey and pink lines connecting regions with direct and inverse sequence identity, respectively. **C)** Zoom-in view of the BLAST alignment of representative isolates in B. ΦESI CDS are colored in blue while non ΦESI CDS are colored in gray. BLAST hits (>99% identity) are represented as grey and pink lines connecting regions with direct and inverse sequence identity, respectively. The associated Arg-tRNA is represented as a green line with a red arrow, and the 1.5 Mbp region between insertion positions is represented as blue and orange box. **D)** Flow diagram describing the distribution of characteristics of the 146 *Salmonella* Infantis complete genomes used in these analyses.

Based on these insertion patterns and their genomic context, the genomes positive for ΦESI sequences were classified into four insertion profiles: profile A (12/105, 11.4%), profile B (64/105, 61.0%), profile C (25/105, 23.8%), and others, assigned to profile D (4/105, 3.8%) (**Table 1**, **Fig. 4C**). In most profile A genomes (66.7%, 8/12) the complete prophage localized to the 1.45 Mbp locus accompanied by partial/fragmentary hits (3.7–7.8 kb) at the 3.05 Mbp locus, exemplified by strain PM57. A reciprocal arrangement, resulting from the inversion, was observed for the remaining profile A genomes (e.g., FSIS12214921), in which ΦESI mapped to the ∼3.05 Mb locus, while fragments mapped to ∼1.45 Mb. In profile B genomes, ΦESI sequences were present in both genomic regions (∼1.45 and ∼3.05 Mb loci), including medium-sized (≈16–36 kb) and small fragments (∼3.7 kb) (e.g., N55391 and Z1324PSL0001). All profile B genomes contained all ΦESI CDSs, and 42.2% (27/64) exhibited the chromosomal inversion. Finally, all profile C genomes carried the chromosomal inversion of the intermediate region and had small (∼3.7 kb) and medium-sized (∼35.8 kb) ΦESI fragments located only at the ≈1.45 Mb locus (e.g., CFSAN024778). Of note, we observed evidence of phylogeographic structuring in the presence and profiles of ΦESI/ΦESI-like prophages in the analyzed genomic datasets. Analysis of 20,429 genomes representative of the global geographic, temporal, niche, and genomic diversity of *Salmonella* Infantis (Enterobase; September 9^th^, 2025), revealed that 94.1% (7,018/7,460) of the ΦESI-positive genomes came from the Americas (**Fig. 3D**). In accordance, analysis of the complete chromosomes revealed that profile B (ΦESI-positive) genomes were enriched among isolates from the Americas (Fisher’s exact test, p=0.014), and profile C (ΦESI-like) genomes were associated with Europe (Fisher’s exact test, p<0.0001) (**Fig. 4D**).

**Table 1.**
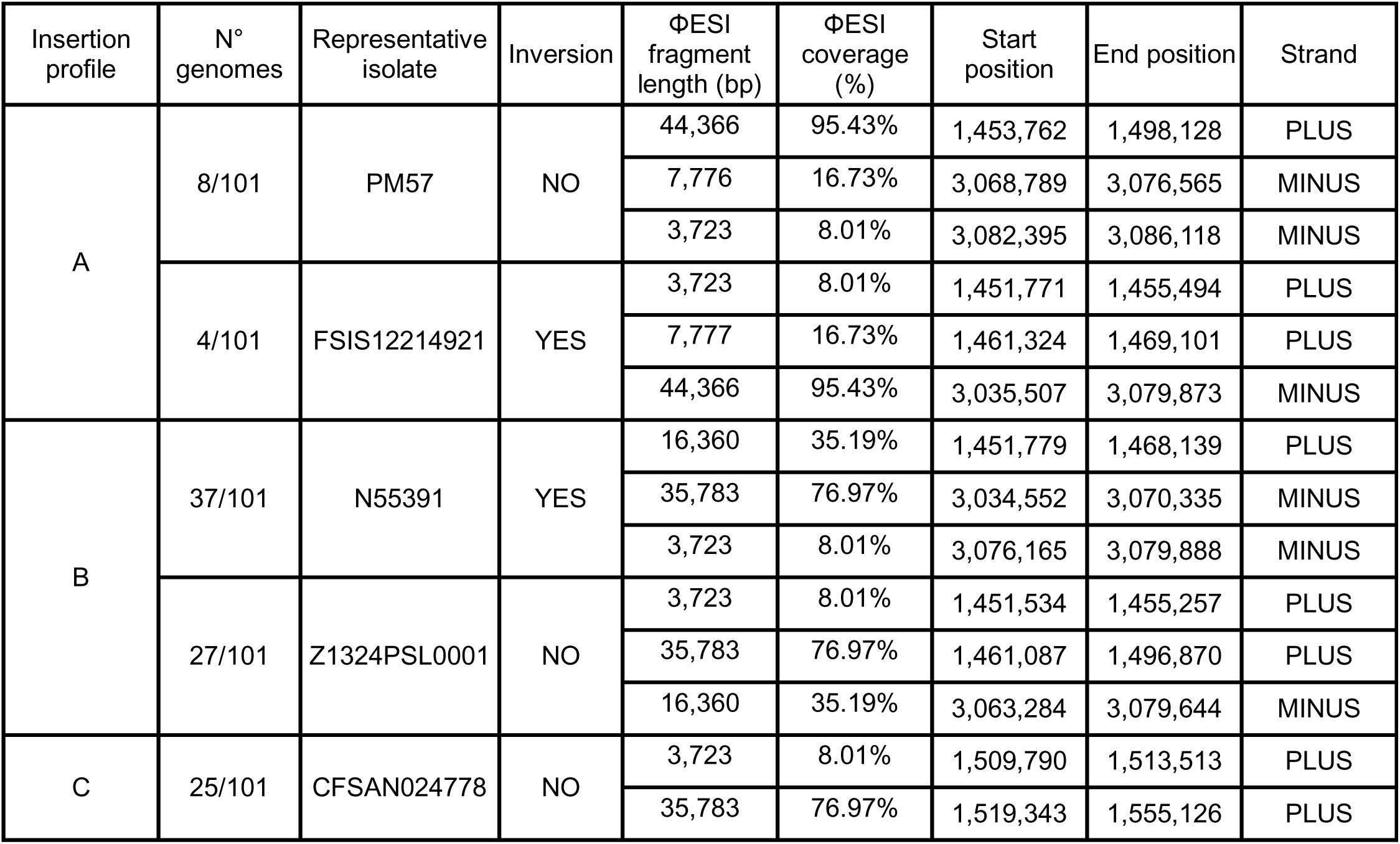
ΦESI insertion profiles among 101 closed genomes of *Salmonella* Infantis.

As ΦESI/ΦESI-like prophages were almost exclusively carried by the emergent *Salmonella* Infantis lineage, and ΦESI from strain PM57 showed lytic activity against the non-emergent DR006 strain, we hypothesized that ΦESI/ΦESI-like phages may provide a competitive fitness to their hosts over ΦESI/ΦESI-like-negative strains.

### **Φ**ESI provides a competitive advantage to emergent *Salmonella* Infantis

Competition assays between PM57 (ΦESI-positive) and DR006 (ΦESI-negative) strains were carried out (**Table S7**). When grown in monocultures, both strains displayed similar growth curves, with no significant statistical differences in their area under the curve (AUC) (**Fig. 5A, C**). However, in coculture, both strains grew following a similar pattern as in monocultures until hour 4, followed by a reduction in the growth of strain DR006 which resulted in a difference of 0.63 log at hour 5, 0.82 log at hour 6, and 0.96 log at 24h (**Fig. 5B**). Accordingly, the AUC representing the growth of DR006 was significantly lower than the AUC for PM57 in coculture, and the AUC of DR006 in monoculture (**Fig. 5C**; p<0.001). The competitive index (CI) of PM57 relative to DR006 in coculture indicated that strain PM57 consistently exhibited higher fitness, as the CI increased above 0 over time despite both strains being inoculated at equal starting concentrations (**Fig. 5E**). This competitive advantage of strain PM57 became even more evident when both strains were quantified directly by strain–specific qPCR (**Fig. S5**). Using this method, the AUC of strain PM57 in coculture remained significantly higher than that of strain DR006, confirming the previous observation (**Fig. 5D**). The CI calculated from absolute qPCR data further demonstrated a clear advantage of strain PM57, with statistically significant CI differences at 4, 5, 6 and 24 h compared with time 0 h (**Fig. 5F**). To quantify ΦESI during the competition assay, PFU/mL estimations on double-layer agar were performed at each time point for both monoculture and coculture samples (**Fig. 5A, B**; blue lines). In monoculture, low PFU levels were detected at 6 and 24 h for PM57, close to the assay’s detection limit (66.6 PFU/mL). No PFUs were detected in the monoculture of strain DR006. In contrast, in coculture, PFUs were detected as early as hour 2 and increased exponentially until hour 6, reaching a maximum of 5.04 x 10^7^ PFU/mL (**Fig. 5B**). The phage in the coculture formed lytic plaques exclusively on DR006 (**Fig. 5G**). These experiments indicate that the release of ΦESI infective particles from strain PM57 negatively impacts the growth of strain DR006, providing a competitive advantage under the coculture conditions.

**Figure 5.**
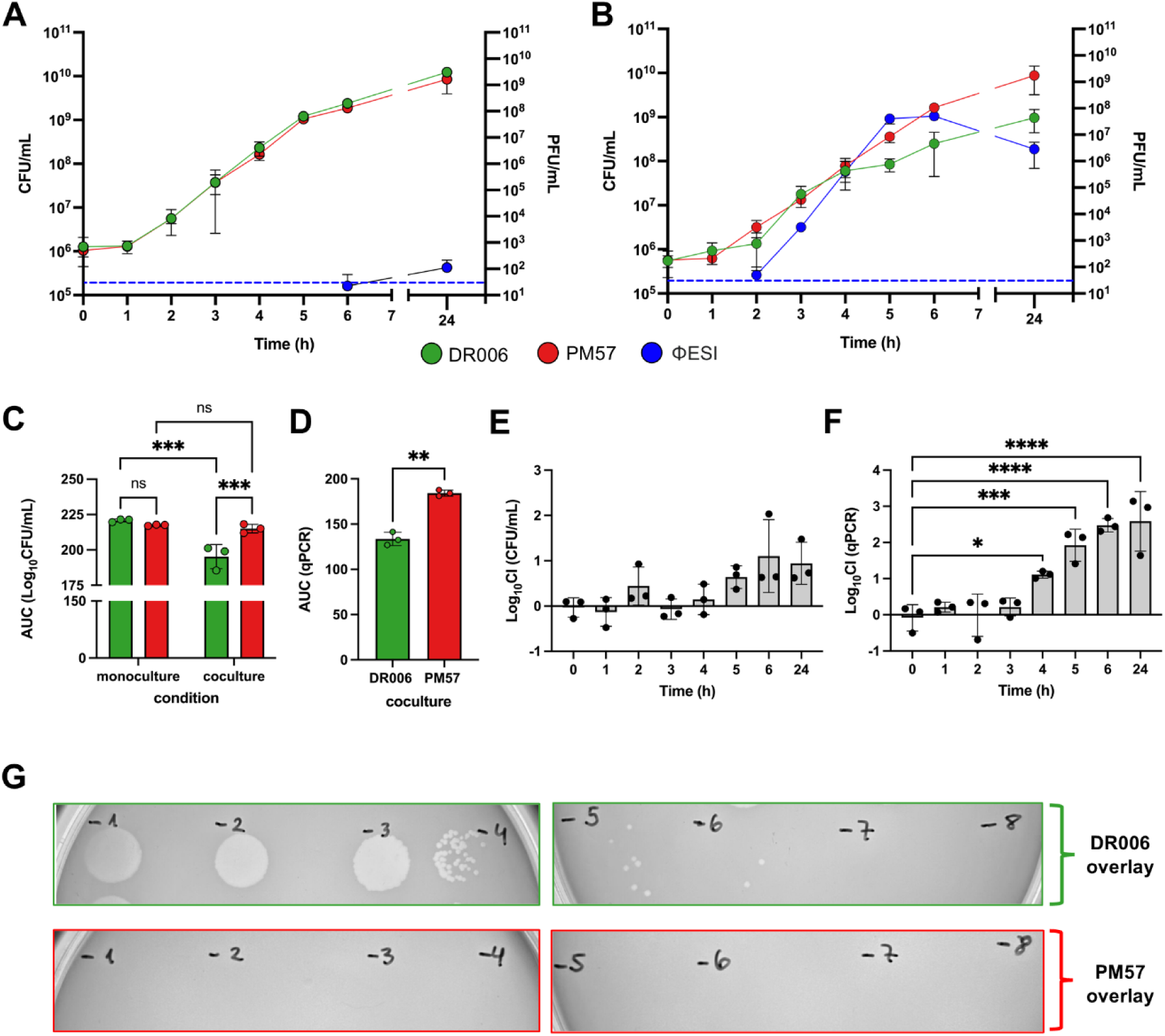
ΦESI-positive *Salmonella* Infantis PM57 has a competitive advantage over isolate DR006 lacking prophage ΦESI. **A)** Growth curves of *Salmonella* Infantis isolates PM57 and DR006 in TSB medium as monocultures or **B)** cocultures. At each time-point the titer of phage particles present in the cultures was determined. The broken blue line represents the limit of detection for phage quantification. The error bars represent the standard deviation of three independent biological replicates. For each time point, quantification were derived from the same experiments, where one aliquot was used for CFU/mL determination (Figure 3B) and a parallel aliquot was used for qPCR-based quantification (Figure S7) **C)** Area under the curve (AUC) of the log10-transformed growth curves shown in A and B. **D)** AUC of the log10-transformed absolute counts of PM57 and DR006 from the experiments in A and B, as determined by qPCR. **E)** The log10 Competitive Index (CI; PM57/DR006), estimated from the experiments in B, is shown based on the CFU titers and **F)** qPCR absolute quantifications. **G)** Sample phage titer quantification showing that lysis plaques formed only when isolate DR006 was used in the agar overlay. See Materials and Methods for statistical details. Statistical significance thresholds were defined as follows: *P* < 0.05 (*), *P* < 0.01 (**), *P* < 0.001 (***), and *P* < 0.0001 (****).

We wanted to assess whether the observed interaction between PM57 and DR006 was not exclusive to these strains. Therefore, efficiency of plating (EOP) assays were carried out with 19 different *Salmonella* Infantis strains representing the diversity of ΦESI/ΦESI-like and pESI megaplasmid carriage (**Table S8**). The presence of ΦESI CDSs in these strains is shown in **Fig. 6A**. We could not find pESI+/ΦESI- nor pESI+/ΦESI-like+ combinations among our sequenced *Salmonella* collection of more than 2,800 isolates. After evaluating the susceptibility of the selected strains, we observed that all 10 ΦESI/ΦESI-like-positive isolates were resistant (EOP=0) to the ΦESI-mediated lysis (**Fig. 6B**). Conversely, most (6/9) of the ΦESI-negative isolates were susceptible (EOP>0.5). Among ΦESI-negative strains evaluated, those with zero ΦESI CDSs showed high values of EOP (about 1.0-1.4). On the other hand, in the strains carrying a few ΦESI CDSs (FA1500, FA1662, FA1547, FA0720) but still ΦESI-negative, the EOP was lower (FA1500, EOP=0.5; 71 CDSs) or zero (FA1662, FA1547, FA0720; 23 CDSs) (**Fig. 6B**), suggesting a possible role for those genes in preventing ΦESI superinfection. Overall, the data indicate that *Salmonella* Infantis carrying ΦESI/ΦESI-like prophages have superinfection resistance while non-carriers remain susceptible.

**Figure 6.**
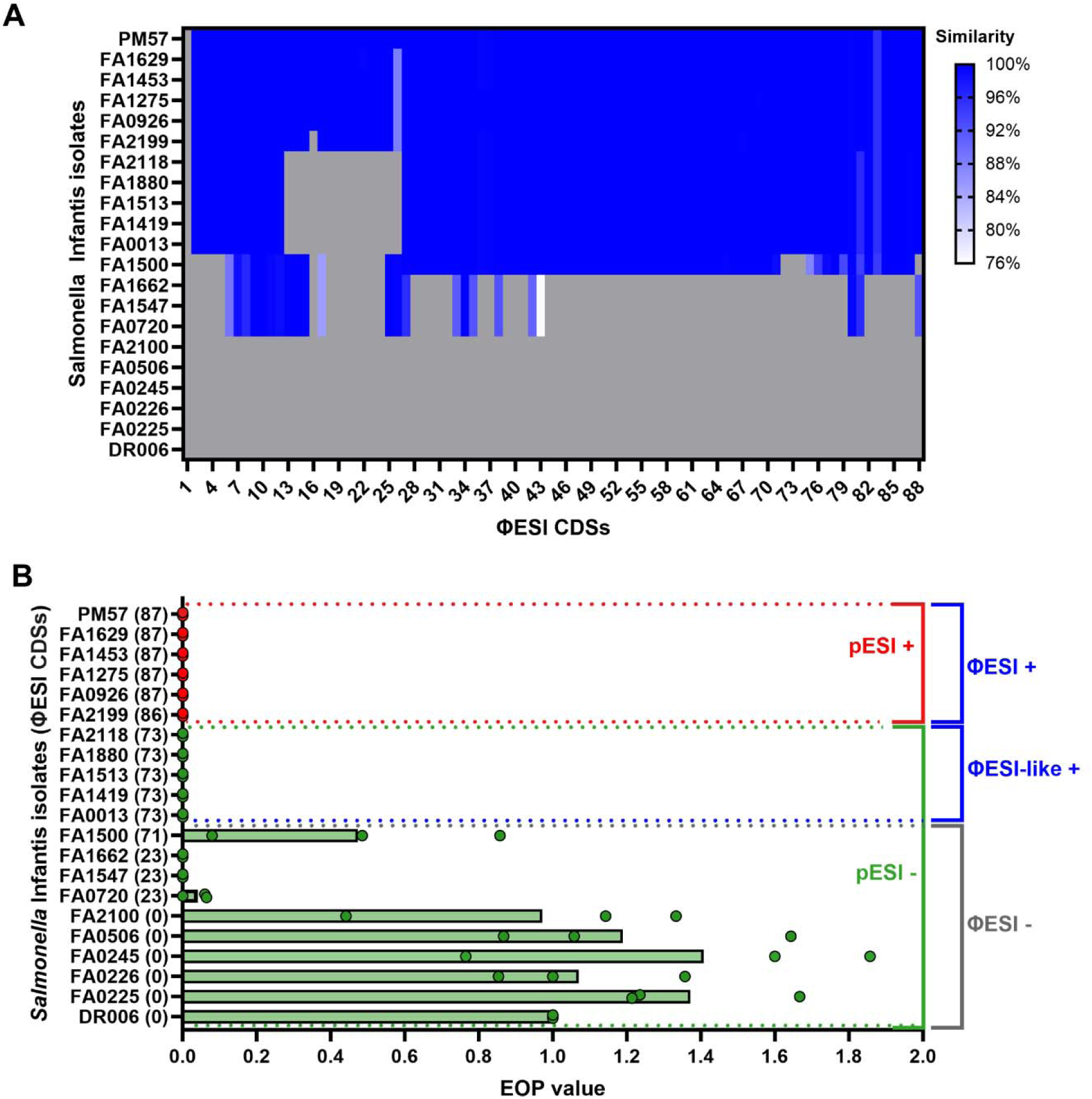
ΦESI and ΦESI-like prophages provides resistance against ΦESI-mediated lysis. **A)** Heatmap showing the presence of ΦESI CDSs (from white to blue) among 21 *Salmonella* Infantis isolates used in the susceptibility to lysis assay. Gray color indicates the absence of a given CDSs. The scale shows a similarity score based on the nucleotide identity and alignment coverage (See methods for details) **B)** The susceptibility of 21 different pESI-positive/negative and ΦESI/ΦESI-like positive/negative *Salmonella* Infantis isolates towards the ΦESI-mediated lysis. The horizontal bars represent the mean susceptibility expressed as the efficiency of plating using strain DR006 as the reference. The individual points indicate the EOP value for each one of the three biological replicates performed. Strain PM57 was also included as a control. The y axis indicates the different isolates tested and shows, in parentheses, the number of ΦESI CDSs present in each isolate. The red and green bars/points indicate the pESI-positive and pESI-negative isolates, respectively. The red and green brackets indicate the presence and absence of pESI-like megaplasmids. The blue and gray brackets indicate the presence and absence of ΦESI/ΦESI-like prophages.

## DISCUSSION

The global dissemination of multidrug-resistant *Salmonella* Infantis represents a global economic burden and public health threat, further increased by the acquisition of ESBL-encoding genes (7). Carriage of the pESI megaplasmids and their variants, which confer enhanced colonization, persistence, and resistance features (8), has been recognized as the major contributor to the success of this foodborne pathogen, while the potential role of additional factors remained unknown.

Here, we discovered and characterized ΦESI, a prophage integrated into the chromosome of the emergent *Salmonella* Infantis PM57 strain, which can form active viral particles that infect and lyse the ΦESI-negative strain DR006, thereby providing a competitive advantage to its host. Previous research, on different bacterial species, has documented that prophage induction in a bacterial subpopulation can confer a competitive advantage to lysogenic bacteria by releasing infective phage particles that selectively lyse and kill susceptible non-lysogenic competitors, while the lysogenic population remains immune due to a superinfection exclusion mechanism (35, 37, 40, 41). Accordingly, our experiments showed the presence of phage-infective particles in the PM57 monoculture, suggesting a basal release of ΦESI into the culture media. In addition, only the isolates carrying ΦESI CDSs showed resistance against ΦESI, while those lacking ΦESI CDSs were susceptible to the ΦESI-mediated lysis. Of note, isolates FA1662, FA1547, and FA0720 harbored 23 CDSs only, but still showed lysis resistance. Considering the presence/absence pattern of the ΦESI CDSs in the tested isolates and their susceptibility, CDSs 006-012, 017, 027, 033-035, 038, 042, 043, 080, and 081 are likely involved in the superinfection exclusion mechanism, that protects against ΦESI. However, most of these CDSs are predicted to encode hypothetical proteins, structural phage proteins, or products with no obvious immunity function. As the bioinformatic and phylogenomic analysis of the *Salmonella* Infantis genomes at a global scale revealed the presence of ΦESI/ΦESI-like prophages almost exclusively within the emergent *Salmonella* Infantis lineage, our results strongly indicate that the observed interaction between the PM57 and DR006 strains is not limited to these two isolates but represents a general phenomenon, suggesting that ΦESI/ΦESI-like prophages have been involved, as well as the pESI megaplasmid, in the successful dissemination of emergent *Salmonella* Infantis across the world.

The widespread presence of ΦESI/ΦESI-like prophages within the emergent *Salmonella* Infantis lineage may have influenced the host range of the conjugative pESI/pESI-like megaplasmids. Prophages can substantially limit the spread of conjugative plasmids by killing susceptible transconjugant bacteria, thereby shrinking the pool of viable plasmid recipients (42). The current evidence shows that the pESI megaplasmids are restricted to the emergent lineage, in which spreads mostly by clonal expansion, and no other *Salmonella* Infantis lineages carry this genetic element (7, 43) (**Fig. 3D**). Therefore, we hypothesize that the ΦESI/ΦESI-like prophages contribute to limiting the acquisition of the pESI megaplasmid by other *Salmonella* Infantis outside the emergent lineage. At the same time, pESI-like megaplasmids may also have actively contributed to the stabilization of ΦESI/ΦESI-like prophages within emergent *Salmonella* Infantis. The CDS 021 of ΦESI is predicted to encode a LexA-family protein of 250 amino acids with 32.8% amino acid identity to the cI repressor of the Lambda phage, containing a Lambda repressor-like DNA-binding domain from positions 3-95, and a LexA/Signal peptidase domain from positions 138-232 (**Fig. S6**). During the SOS response, the activated RecA protein (RecA*) cleaves the cI repressor between the Ala_111_-Gly_112_ residues, allowing the transcription of the Lambda lytic-associated genes (44). These residues are also conserved in the CDS 021 (**Fig. S6**), suggesting that induction of the ΦESI lytic cycle could be triggered by the host SOS response. The pESI/pESI-like megaplasmids harbor a homolog (locus tag APL92_RS24015 in pN55391) of the *psiB* gene that encodes the RecA-binding protein with anti SOS-response activity, thereby protecting the host from lysis by preventing inadvertent prophage induction during stressful conditions (44–46). Additionally, the loss of this plasmid-encoded safeguard can leave hosts more vulnerable to SOS-driven prophage activation (47). Together, these mechanisms could have favored the coexistence of ΦESI/ΦESI-like prophages and pESI/pESI-like megaplasmids within emergent *Salmonella* Infantis, shaping the successful spread of this pathogen.

Bacterial chromosomes can undergo large inversions, commonly via homologous recombination between repeated DNA sequences and mobile elements, including prophages, are frequent mediators or markers of chromosomal rearrangements (48). ΦESI is integrated at the 3’-end of an Arg-tRNA located approximately at 1.5 Mbp in strain PM57. However, we also detected ΦESI genes at position ≈3.05 Mbp, separated by an intermediate region of ≈1.45 Mbp (**Fig. 2A**). We observed the same overall pattern across 105 out of 146 closed *Salmonella* Infantis chromosomes, displaying 4 different arrangements based on the location of ΦESI at either the ≈1.45 or ≈3.05 Mbp loci, and the surrounding genetic context. We detected that in 39.0% of the chromosomes, a large-scale inversion of the ≈1.5 Mbp intermediate region was present, dramatically changing the genome synteny (**Fig. 4**). While DNA rearrangements may confer adaptive advantages resulting from gene-dosage effects derived from proximity to the replication origin (48, 49), the large-scale inversions that we detected in emergent *Salmonella* Infantis likely do not perturb transcriptional balance or mean gene copy number as they are almost symmetrical to the *oriC*, thereby preserving gene distances.

Our study is not free of limitations, and additional experiments would strengthen the evidence of a direct cause-and-effect relationship between ΦESI and the competitive advantage over ΦESI-negative strains. For this, the construction and testing of a PM57 strain lacking ΦESI would be helpful to assess the role of this prophage alone, enabling competition experiments against a strain with an isogenic background. Also, obtaining DR006 mutants resistant to ΦESI-mediated lysis would allow to identify the ΦESI receptor or phage resistance mechanisms, and would enable testing whether susceptibility to ΦESI is the only DR006-associated factor contributing to its reduced fitness when cocultured with PM57. In addition, curing of pESI from PM57 would allow to directly test whether plasmid carriage alters ΦESI infection dynamics, lysogeny frequency, phage production and the competitive phenotype observed in coculture. Moreover, while our results indicate that ΦESI infective particles are constitutively induced at low basal levels (**Fig. 4A**), we did not investigate the potential role of external factors (e.g. temperature, secreted factors of competing bacteria, antibiotic at subinhibitory concentrations, etc.) as prophage inducers, which may increase the release of infective particles under biologically relevant conditions, contributing to a more complete understanding of the ecological role of ΦESI in the emergence and successful dissemination of emergent *Salmonella* Infantis.

## Conclusions

Pathogen emergence is a complex, multifactorial process in which horizontal gene transfer plays a critical role (50). Our findings provide strong evidence that ΦESI and ΦESI-like prophages are active contributors to the ecological and adaptative dynamics of the emergent *Salmonella* Infantis lineage by functioning as a competitive weapon able to selectively eliminate susceptible competitors and potentially shaping the distribution of the pESI/pESI-like megaplasmids. The coexistence of both mobile elements within emergent *Salmonella* Infantis likely involves still-unexplored interactions that may have contributed to the stabilization, persistence, and global dissemination of this successful lineage.

## MATERIALS AND METHODS

### Bacterial strains and growth conditions

Twenty-one *Salmonella* Infantis isolates isolated from surface waters (n=19) (18), poultry meat (n=1) (15), and poultry raised in backyard production systems (n=1) (51) were used for experiments in this work. Detailed information on these isolates is provided in **Table S8** of the supplementary material. *Salmonella* Infantis was cultured in Trypticase Soy Broth (TSB) or Trypticase Soy Agar (TSA; agar 1.5% w/v) at 37 °C. Tetracycline (20 µg/mL) was added to TSA plates when required.

### Phage isolation and purification

Phage ΦESI was isolated from a coculture of the two *S*I strains DR006 and PM57. One hundred mL of TSB was inoculated with 500 µL of the overnight culture of each *Salmonella* Infantis strain and incubated at 37 °C overnight. One mL of the coculture was filtered through a 0.22 µm pore-size syringe filter. Then, 10 µL of the filtrate was spotted on a double-layer agar of the host (PM57 or DR006). After overnight incubation at 37°C, the presence or absence of lysis was observed. Plaques were further purified by 4 passages. Lysis (for the first passage) or single plaques (for the second and third passages) were purified by picking with a micropipette tip into the plaque, followed by resuspending the phages in SM buffer (100 mM NaCl, 8 mM MgSO_4,_ 50 mM Tris-HCl, pH 7.5). A single plaque of the fourth passage was stung and used for phage amplification and preparation of a concentrated phage solution (lysate) by precipitation using PEG8000 (Polyethylene Glycol 8000) for further experiments.

### Transmission Electron Microscopy

Morphotype of ΦESI was determined by transmission electron microscopy (TEM). Twelve µL of the concentrated phage solution (10^10^ PFU/mL) was stained with 2% uranyl acetate and then added to a copper TEM grid. The grid with the stained sample was observed using a transmission electron microscope at the Advanced Microscopy Unit at Pontificia Universidad Católica de Chile (Talos^TM^ F200X G2 TEM, Detection Ceta 16M CMOS Camera, pixel size 14 µm x 14 µm, 16 bit; Electron source: X-FEG Field Emission Gun). The images were analyzed using Fiji ImageJ (version: 2.14.0/1.54f).

### DNA extractions and purification

Phage genomic DNA was extracted from a concentrated phage lysate (10¹ PFU/mL) using phenol–chloroform extraction followed by ethanol precipitation, as previously described (52), with minor modifications. Briefly, 1 mL of concentrated phage suspension was treated with 5 µL of DNase I (1 µg/mL), 6 µL of RNase A (30 µg/mL), and 10 µL of CaCl_2_ (100 mM). This mix was incubated at room temperature for 30 min to remove contaminating host nucleic acids. DNase I activity was subsequently inactivated by the addition of 40 µL of EDTA (0.5 M). Protein digestion was performed by adding 6 µL of proteinase K (20 mg/mL) and 30 µL of SDS (20%, w/v), then incubation at 56 °C for 1 h. After cooling to room temperature, an equal volume (1 mL) of phenol was added, mixed gently by inversion, and centrifuged at 2,356g (5,000 rpm) for 5 min. The aqueous phase was carefully transferred to a new tube and extracted with an equal volume of phenol:chloroform (1:1), then centrifuged under the same conditions. The aqueous phase was transferred to a fresh tube and further extracted with an equal volume of chloroform, then centrifuged as described above. DNA was precipitated by adding 0.1 volumes of sodium acetate (3 M), followed by the addition of two volumes of 100% ethanol. Samples were gently mixed and incubated overnight at 4 °C. DNA was pelleted by centrifugation at 14,000g (12,000 rpm) for 15 min at 4 °C. The supernatant was discarded, and the pellet was washed with 100% ethanol, followed by centrifugation under the same conditions. After carefully removing the supernatant, the pellet was air-dried at room temperature until residual ethanol had completely evaporated. Finally, the DNA was resuspended in 50 µL of nuclease-free water.

Genomic DNA (gDNA) of *Salmonella* Infantis PM57 was purified from 3 mL of an overnight culture grown in TSB at 37°C. The bacterial culture was pelleted by centrifugation at 8000g (9,069 RPM) during 7 min. at 4°C and then resuspended in 180 μL of buffer ATL from the QIAGEN DNeasy Blood & Tissue kit. Twenty microliters of Proteinase K (600 mAU/mL) were added and mixed by vortexing. The suspension was incubated at 56°C for 30 min for cell lysis. Total DNA was extracted using phenol:chloroform:isoamil alcohol, precipitated with 3M sodium acetate (pH5.2) and 100% propan-2-ol, washed with 75% ethanol, and resuspended in 50 μL of nuclease-free water, as previously described (53). Genomic DNA from the competition assays was extracted using the same procedure outlined for PM57.

The concentration and purity of the obtained phage and bacterial DNA were assessed using a Nanodrop spectrophotometer or a Qubit™ 3.0 fluorometer. The DNA was stored at -20°C until required.

### Whole genome sequencing (WGS), assembly and annotation

Bacteriophage genomic DNA was sequenced using Illumina technology. DNA samples were submitted to MicrobesNG (Birmingham, UK) for library preparation and sequencing. Genomic libraries were prepared using the Nextera XT Library Prep Kit, following the manufacturer’s protocol with minor modifications: the input DNA amount was increased two-fold, and the PCR elongation time was extended to 45s to improve library complexity. DNA quantification and library preparation were performed using a Hamilton Microlab STAR automated liquid handling system. Libraries were sequenced using a 250 bp paired-end protocol. Raw reads were quality-filtered and adapter-trimmed using Trimmomatic v0.30 (54) with a sliding-window quality cutoff of Q15. *De novo* genome assembly was performed using SPAdes v3.7 (55). Contigs were annotated using Pharokka v1.7.5 with the v1.4.0 database (56).

Purified gDNA of *Salmonella* Infantis PM57 was sent to SeqCenter LLC (Pittsburgh, USA) for hybrid WGS using Illumina and Oxford Nanopore Technologies (ONT) platforms, and genome assembly. Illumina sequencing libraries were prepared using the tagmentation-based and PCR-based Illumina DNA Prep kit and custom IDT 10bp unique dual indices (UDI) with a target insert size of 280 bp. No additional DNA fragmentation or size selection steps were performed. Illumina sequencing was performed on an Illumina NovaSeq X Plus sequencer in one or more multiplexed shared-flow-cell runs, producing 2x151bp paired-end reads. Demultiplexing, quality control, and adapter trimming were performed with bcl-convert (v4.2.4). Long-read libraries were prepared using the PCR-free ONT Ligation Sequencing Kit (SQK-NBD114.96) with the NEBNext® Companion Module (E7180L) according to manufacturer’s specifications. No additional DNA fragmentation or size selection was performed. Nanopore sequencing was performed on an ONT GridION or PromethION sequencer using R10.4.1 flow cells in one or more multiplexed shared-flow-cell runs. The run design used the 400 bps sequencing mode with a minimum read length of 200 bp. The Dorado v0.5.3 basecaller (https://github.com/nanoporetech/dorado) was used under the super-accurate (sup) and 5mC/5hmC (5mC_5hmC) basecalling models. Dorado demultiplex was used under the SQK-NBD114.96 kit settings to generate the BAM file. SAMtools fastq v1.20 (57) was used to generate the FASTQ file from the BAM file.

Porechop v0.2.4 (https://github.com/rrwick/Porechop) was used with default parameters to trim residual adapter sequences from the ONT reads that may have been missed during basecalling and demultiplexing. *De novo* genome assemblies were generated from the ONT read data with Flye v2.9.2 (58) under the nano-hq (ONT high-quality reads) model. Disjointing assemblies with the longest reads that provided 50x coverage (--asm-coverage), assuming a genome size of 6 megabases (--genome-size), were included. Subsequent polishing used the Illumina read data with Pilon v1.24 (59) under default parameters. To reduce erroneous assembly artifacts caused by low quality nanopore reads, long read contigs with an average short read coverage of 15x or less were removed from the assembly. Assembled contigs were evaluated for circularization via Circlator v1.5.5 (60) using the ONT long reads. Assembly annotation was performed with Bakta v1.10.1 using the Bakta v5.1 database (61).

### Taxonomic analysis

To assess the taxonomy of ΦESI, a BLASTn search against the NCBI Viral Nucleotide Collection in NCBI Virus (https://www.ncbi.nlm.nih.gov/labs/virus/vssi/#/) was performed using the ΦESI assembly as query. The genomes from the top-eight BLASTn results were downloaded and used to carry out taxonomic analyses using the VICTOR and VIRIDIC web platforms, using default parameters and the d6 equation for VICTOR (62, 63).

### Distribution analysis of **Φ**ESI genes among global *Salmonella* Infantis genomes

Metadata, assembly statistics, and cgMLST-based hierarchical clusters information were retrieved for all 36,499 *Salmonella* Infantis genomes publicly available in Enterobase (64) as of September 9^th^, 2025. Low-quality and redundant genomes were filtered out as previously described (7). Briefly, assemblies with N50<100,000 and coverage<30X were removed from the dataset. Redundant genomes with the same collection year, source niche, country of origin, and from the same HC5 cluster (linked by ≤5 cgMLST allele differences) were removed leaving only one representant genome in order to retain most of the temporal, geographic, isolation source and genomic diversity. After this process, 20,429 genomes were selected and downloaded for further analysis (**Table S2**).

The presence of the ΦESI genes in the global *Salmonella* Infantis dataset was assessed using ABRicate v1.0.1 (https://github.com/tseemann/abricate) and a custom database made with the 88 ΦESI CDSs identified with Pharokka. After examination of the gene-number distribution across all 20,429 *Salmonella* Infantis genomes, we considered the presence of 87 ΦESI CDS as indicative of a complete ΦESI prophage, 73-86 ΦESI CDSs as a ΦESI-like prophage, and <73 ΦESI CDSs as negative for ΦESI (**Table S3**). The presence of pESI-like megaplasmids was also assessed with ABRicate and a custom database as previously described (7).

A recombination-free core-SNP maximum likelihood phylogenomic tree was constructed based on 864 non-redundant *Salmonella* Infantis genomes (according to collection year, country, and HC50 information) from the cluster HC200_36, which encompasses all genomes positive for ΦESI and ΦESI-like prophages. This phylogeny was constructed as recently described (7). Briefly, the whole genome alignment was obtained with Snippy v.4.6.0 (https://github.com/tseemann/snippy) using the contigs of the 864 genomes and the closed chromosome of *Salmonella* Infantis strain N55391 as the reference (Enterobase assembly barcode SAL_XA0374AA_AS), using default parameters. Recombination regions were detected with Gubbins v3.3 (65) after 5 iterations using the iqtree-fast option and the Jukes-Cantor (JC) model for the first iteration, and the –best-model option for the remaining iterations. The whole-genome alignment with masked recombination regions obtained from Gubbins was used as the input to snp-sites v2.5.1 (66) to obtain the recombination-free core-SNP alignment. The core-SNP phylogenomic tree was constructed with IQ-Tree MPI multicore v2.3.2 (67) using the GTR+G4+F+ASC nucleotide substitution model, which was selected in the Gubbins iterations. Node support was assessed with 1,000 ultrafast bootstraps (68). The resulting tree was uploaded to iTOL v7 (69) and rooted to the best-fitting temporal root estimated with TempEst v1.5.3 (70). The geographic information (Regions of origin; geoscheme of the United Nations), megaplasmid presence, and number of ΦESI CDSs were integrated to the phylogeny.

### Analysis of prophage **Φ**ESI genomic context

A new custom ABRicate database, containing one sequence corresponding to the complete ΦESI genome was used to screen all available *Salmonella* Infantis genomes retrieved from NCBI Pathogen Detection (May 22^nd^, 2025); only genomes designated as “Complete” in Pathogen Detection were included in the screening (**Table S4**). ABRIcate summary outputs were used to define presence/absence matrices and to identify distinct prophage insertion profiles across the dataset.

For each identified insertion site, the genomic context was defined by extracting 10 kb upstream and 10 kb downstream of the insertion coordinates from representative genomes. Flanking sequences were clustered with cd-hit-est v4.8.1 (https://github.com/weizhongli/cdhit) at 99% nucleotide identity, to reduce redundancy and to group highly similar genomic contexts; a single representative sequence from each cluster was selected for downstream visualization and comparative interpretation. Custom scripts were used to parse ABRicate outputs, extract flanking regions, perform clustering, and select representative genomes/contexts. These scripts are available in the project repository (see Data Availability). Finally, representative genomes for each unique insertion profile (and separately for genomes exhibiting large-scale structural rearrangements) were chosen for whole-genome structural comparison and visualization using pyGenomeViz (https://github.com/moshi4/pyGenomeViz). All bioinformatic analyses were performed on a Linux computer server using 12 CPU threads unless otherwise stated; default tool parameters were used unless specified above.

### Competition assay between emergent and non-emergent *Salmonella* Infantis

A competition assay was performed between *Salmonella* Infantis strain PM57 (emergent clade) and strain DR006 (non-emergent). Individual, overnight cultures of the two strains were prepared in TSB at 37°C. Both strains were cocultured in 100 mL TSB at an initial concentration of 10 CFU/mL (1:1 ratio) and incubated at 37°C and 175 rpm. Samples were collected hourly for six hours and again after 24 hours to quantify phage concentration (PFU/mL) in a double-agar overlay, and bacterial load by colony counting (CFU/mL).

To estimate phage concentration, aliquots (<1 mL) from the cocultures were filtered through a 0.22 µm pore-size syringe filter. Subsequently, 20 µL of the filtrate was serially diluted tenfold (up to 10) in SM buffer. Then, 5 µL of each dilution was spotted in triplicate onto double-layer agar plates containing *Salmonella* Infantis DR006 as the host strain. Then, the plates were incubated overnight at 37 °C. Plaque-forming units were counted to determine the concentration expressed in PFU/mL. All assays were performed independently three times (three biological replicates).

To estimate the total bacterial loads (PM57+DR006), 20 µL of each sample was serially diluted tenfold (up to 10) in 0.9% NaCl. Then, 10 µL of each dilution was spotted in triplicate onto TSA plates and incubated at 37°C overnight. To specifically quantify strain PM57, 10 µL of each dilution was also spotted in triplicate onto TSA supplemented with tetracycline (20 µg/mL), as PM57 is resistant to tetracycline (IC_50_=163.9 µg/mL) while DR006 is susceptible (IC_50_=1.5 µg/mL) (**Fig. S7**). Plates were incubated overnight at 37 °C.

The concentration of strain DR006 was calculated indirectly by subtracting the obtained value for PM57 from the total bacterial load according to the following formula:

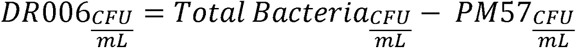

As controls, each strain was also cultured individually (monoculture) under the same conditions, and bacterial concentrations were determined at the same time points.

Competitive Index (CI) of PM57 over DR006 in the coculture at each time point was calculated according to the following formula:

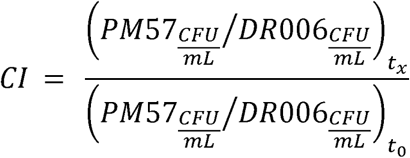

The denominator of the formula represents the average 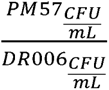 ratio at time 0, calculated from three independent biological replicates.

### qPCR quantification

Targeted quantitative PCR (qPCR) assays were performed to distinguish between PM57 and DR006 in the competition assays and to estimate their absolute loads (**Fig. S8, S9**). For PM57 quantification, qPCR reactions were conducted using 50 ng of template DNA per reaction and primers targeting the baseplate X gene, which is present in the PM57 chromosome but absent from the DR006 chromosome (**Fig. S8**). The following primers were used: PM57_qPCR_Fw 5’-CTGAACGTCGGGCAGTT-3’ and PM57_qPCR_Rev 5’- CTGAGGGCGTGGTTGAG-3’, which produce an amplicon of 101 bp in strain PM57 but not in DR006. For DR006 detection, qPCR reactions were carried out using 50 ng of template DNA per reaction and primers targeting a genomic region spanning the interval between the *fieF* and *cpxP* genes. In the PM57 chromosome, these primers are separated by 31,676 bp (**Fig. S8**); therefore, no amplification is expected in PM57 under standard qPCR conditions, which cannot amplify such a large genomic fragment, ensuring assay specificity for DR006. The primers used were: DR006_qPCR_Fw 5’-GCTAATAACAGAATGACAAACTGGAG-3’ and DR006_qPCR_Rev 5’-CAACTCACGTTCCCAGTAACA-3’, producing an amplicon of 113 bp in DR006.

All qPCR reactions were performed with the PowerUp™ SYBR™ Green Master Mix (Applied Biosystems) in a QuantStudio^TM^ 5 real-time PCR system with the QuantStudio^TM^ Design & Analysis Software v1.5.2, following the master mix specifications.

### Susceptibility assays on *Salmonella* Infantis strains carrying **Φ**ESI and **Φ**ESI-like prophages

The susceptibility to the lysis mediated by ΦESI viral particles of different *Salmonella* Infantis representing pESI+/ΦESI+, pESI-/ΦESI-like+, and pESI-/ΦESI-isolates (**Table S1**) was determined using the efficiency of plating (EOP) assay. A suspension of phage ΦESI was serially diluted from 10^-1^ to 10^-8^ in SM buffer, and 5 µL of each dilution was spotted on a soft TSA agar overlay (0.7% agar w/v) seeded with each *Salmonella* isolate. The plates were incubated overnight at 37°C. The efficiency of plating was calculated by dividing the resulting phage titer of the tested isolate by the titer of the reference isolate *Salmonella* Infantis DR006.

### Statistical analysis

Statistical analyses were performed to compare growth dynamics and competitive performance between bacterial strains (PM57 and DR006 in competition). Growth curves were analyzed by calculating the area under the curve (AUC). Differences in AUC between monocultures and cocultures for each strain were assessed using a two-way ANOVA followed by multiple comparison tests. For AUC values obtained from qPCR-based quantification, comparisons were performed using Welch’s t-test to account for unequal variances between strains in coculture. Competition index values across time points were analyzed using an ordinary one-way ANOVA followed by Dunnett’s multiple comparisons test, where all time points were compared against time 0 as the control condition. Statistical significance was defined according to the indicate *P values* for each statistical analysis. All statistical analyses were performed using GraphPad Prism Version 10.2.3.

The association between the presence of ΦESI/ΦESI-like prophages and global emergent *Salmonella* Infantis lineage (represented by carriage of pESI-like megaplasmids) (7) was assessed with a χ^2^ test. 95% confidence interval (CI) computed on the log scale using the Wald approximation. The geographic distribution of insertion profiles of ΦESI/ΦESI-like prophages among *Salmonella* Infantis isolates with an available complete genome was assessed through two-sided Fisher’s exact tests. All tests were performed in GraphPad Prism Version 10.2.3, with α=0.05.

## Funding

Powered@NLHPC: This research was partially supported by the supercomputing infrastructure of the NLHPC (CCSS210001).

Agencia Nacional de Investigación y Desarrollo de Chile (ANID), Fondecyt Postdoctorado 3230796 (API), Fondecyt Regular 1231082 (AIMS), Fondecyt Iniciación 11250702 (DAE), Beca de Doctorado Nacional Folio 21220654 (RBM), Beca de Doctorado Nacional Folio 21241909 (DTN).

## Data availability

All data is available in this manuscript and its Supplemental Materials. Genome assemblies of ΦESI and PM57 are available at NCBI under accession numbers <u>PZ014354.1</u> and <u>JBUZKO000000000.1</u>, respectively.

All custom scripts used in this study are available at the GitHub repository <u>BaFoP-Lab/phiESI_2026_code_repo</u>.

## CRediT authorship contribution statement

**CMP**: Data curation, Formal analysis, Investigation, Methodology, Validation, Visualization, Writing – original draft, Writing – review & editing; **DTN**: Data curation, Formal analysis, Investigation, Validation, Visualization, Writing – original draft, Writing – review & editing; **RBM**: Formal analysis, Investigation, Methodology, Writing – original draft, Writing – review & editing; **IHV**: Investigation, Writing – original draft, Writing – review & editing; **JAU**: Writing – original draft, Writing – review & editing; **DAE**: Formal analysis, Investigation, Methodology, Resources, Writing – original draft, Writing – review & editing; **AIMS**: Funding Acquisition, Methodology, Resources, Supervision, Writing – original draft, Writing – review & editing; **API**: Conceptualization, Data curation, Formal analysis, Funding Acquisition, Investigation, Methodology, Resources, Validation, Supervision, Visualization, Writing – original draft, Writing – review & editing.

